# A genome-wide screen reveals that Dyrk1A kinase promotes nucleotide excision repair by preventing aberrant co-stabilization of cyclin D1 and p21

**DOI:** 10.1101/2022.04.14.488378

**Authors:** François Bélanger, Cassandra Roussel, Christina Sawchyn, Sari Gezzar-Dandashi, Aimé Boris Kimenyi Ishimwe, Frédérick Antoine Mallette, Hugo Wurtele, Elliot Drobetsky

## Abstract

Nucleotide excision repair (NER) eliminates highly-genotoxic solar UV-induced DNA photoproducts that otherwise stimulate malignant melanoma development. Here, a genome-wide loss-of-function screen, coupling CRISPR/Cas9 technology with a flow cytometry-based DNA repair assay, was used to identify novel genes required for efficient NER in primary human fibroblasts. Interestingly, the screen revealed multiple genes encoding proteins, with no previously known involvement in UV damage repair, that significantly modulate NER uniquely during S phase of the cell cycle. Among these, we further characterized Dyrk1A, a dual specificity kinase that phosphorylates the proto-oncoprotein cyclin D1 on threonine 286 (T286), thereby stimulating its timely cytoplasmic relocalization and proteasomal degradation required for proper regulation of the G1-S phase transition and control of cellular proliferation. We demonstrate that in UV-irradiated HeLa cells, depletion of Dyrk1A leading to overexpression of cyclin D1 causes inhibition of NER uniquely during S phase and reduced cell survival. Consistently, expression/nuclear accumulation of nonphosphorylatable cyclin D1 (T286A) in melanoma cells strongly interferes with S phase NER and enhances cytotoxicity post-UV. Moreover, the negative impact of cyclin D1 (T286A) overexpression on repair is independent of cyclin-dependent kinase activity but requires cyclin D1-dependent co-stabilization of p21. Our data indicate that inhibition of NER during S phase might represent a previously unappreciated non-canonical mechanism by which oncogenic cyclin D1 fosters melanomagenesis.

## INTRODUCTION

Nucleotide excision repair (NER) is the preeminent pathway in humans for removing helix-destabilizing, replication and transcription-blocking DNA adducts generated by a variety of environmental mutagens/carcinogens and chemotherapeutic drugs. Among such adducts are solar UV-induced cyclobutane pyrimidine dimers and 6-4 pyrimidine-pyrimidone photoproducts (6-4PP), which represent the overriding cause of mutations promoting cutaneous tumour development(1, 2). NER comprises two overlapping sub-pathways, differing only in the mechanism of DNA damage recognition. During <u>G</u>lobal <u>G</u>enomic <u>NER</u> (GG-NER), helix-distorting DNA lesions are recognized throughout the genome by heterotrimeric XPC/HR23B/Centrin2, in collaboration with the DDB2/DDB1/Cul4A ubiquitin E3 ligase complex. The other sub-pathway, transcription-coupled NER, acting only along the transcribed strand of active genes, is initiated when elongating RNA polymerase II stalls at damaged DNA bases, which in turn promotes recruitment of CSB and the CSA/DDB1/Cul4A ubiquitin E3 ligase complex. After lesion recognition associated with either sub-pathway, proteins of the “core NER pathway” are sequentially recruited and function as follows: (i) The helicase and ATPase activities of XPD and XPB, respectively, as subunits of the TFIIH basal transcription factor, mediate DNA unwinding at the damaged site; (ii) XPA partners with heterotrimeric replication protein A (RPA) to stabilize the unwound DNA and promote lesion verification in collaboration with TFIIH; (iii) The ERCC1-XPF and XPG endonucleases incise the DNA backbone on either side (5’ and 3’ respectively) of the lesion, producing a single-stranded DNA fragment containing the adduct which is subsequently excised; (iv) the resulting ∼30 bp gap is resynthesized by DNA replication factors using the damage-free complementary strand as template and, finally, (v) DNA ligases seal the remaining nick to restore the original DNA sequence. (For review of the NER pathway see(3, 4)).

The syndrome *Xeroderma pigmentosum* (XP), characterized by homozygous germline mutations in NER pathway genes and remarkable (up to 5000-fold increased) susceptibility to sunlight-induced skin cancers including malignant melanoma (MM), underscores the importance of NER to human health(5). In view of this, it may be surprising that multiple genome-wide sequencing studies have revealed a paucity of NER gene defects in sporadic MM within the general population(2, 6, 7). On the other hand, such studies have identified numerous potential MM driver mutations, although in many cases it remains unclear how these mutations promote melanoma development, and whether some might do so by negatively impacting NER.

We previously demonstrated that defective responses to DNA replication stress, e.g. in cells depleted for ATR kinase or translesion DNA polymerase eta (pol eta), cause GG-NER defects during S phase, whereas repair in G1 or G2/M is not significantly impacted (GG-NER during S is hereafter denoted <u>S P</u>hase <u>R</u>epair; SPR)(8–11). As outlined above, the RPA complex, which binds and protects single-stranded DNA (ssDNA) generated during genotoxin-induced replicative stress (12), also plays a critical role in NER(13). Upon severe UV-induced replication stress, excessive sequestration of RPA on ssDNA located at persistently-stalled replication forks, or at aberrantly activated origins of replication, was shown by our group and others to limit the availability of this complex to act in NER during S phase (10, 11, 14). This raises the possibility that a multitude of proteins which act to mitigate replicative stress, and hence ssDNA generation, might modulate the efficiency of NER in an S phase-specific manner. Moreover, such proteins may have thus far escaped detection because classical NER assays, e.g., quantification of NER gapfilling by unscheduled DNA synthesis (UDS)(15), are not designed to evaluate UV damage repair specifically during S.

Here, we employed a FACS-based CRISPR/CAS9 genome-wide screen to identify factors required for efficient removal of UV-induced DNA photoproducts in primary human fibroblasts. Interestingly, this screen revealed multiple proteins that significantly influence NER uniquely during S phase, including Dyrk1A kinase which phosphorylates the proto-oncoprotein cyclin D1 on threonine 286 (T286)(16). Failure to modify cyclin D1 on T286 leads to aberrant nuclear accumulation of the protein, premature S phase entry, and enhanced cellular proliferation that drives tumourigenesis(17). We demonstrate here that lack of Dyrk1A inhibits S phase NER and cell survival post-UV by causing overexpression of cyclin D1. Moreover, this does not depend on either modulation of cyclin-dependent kinase (CDK) activity or the induction of DNA replication stress, but instead on cyclin D1-dependent upregulation of p21 expression.

## RESULTS

### A CRISPR/Cas9 screen identifies novel genes regulating NER

We previously developed a flow cytometry-based immunoassay to directly quantify the repair of 6-4PP as a function of cell cycle(9). Here, this assay was exploited in conjunction with CRISPR/Cas9 technology to perform a genome-wide loss of function screen as a means of identifying genes that promote efficient NER. LF-1 primary lung fibroblasts(18) were transduced at low multiplicity of infection with pool A of the GeCKO v2 single-vector system(19), followed by 10 days of puromycin selection to remove uninfected cells. 10^8^ selected cells were then irradiated with 20 J/m^2^ UV, and ones presenting high residual 6-4PP at 5h post-irradiation were sorted by flow cytometry (Figure 1A). Although the harsh conditions of our labeling protocol adversely affected the yield of intact single cells, we successfully sorted 1.2×10^7^ individual cells resulting in a coverage of approximately 200 genome equivalents. A sample of 30 million unsorted cells was used as control to assess the sgRNA content of the transduced cell population.

**FIGURE 1:**
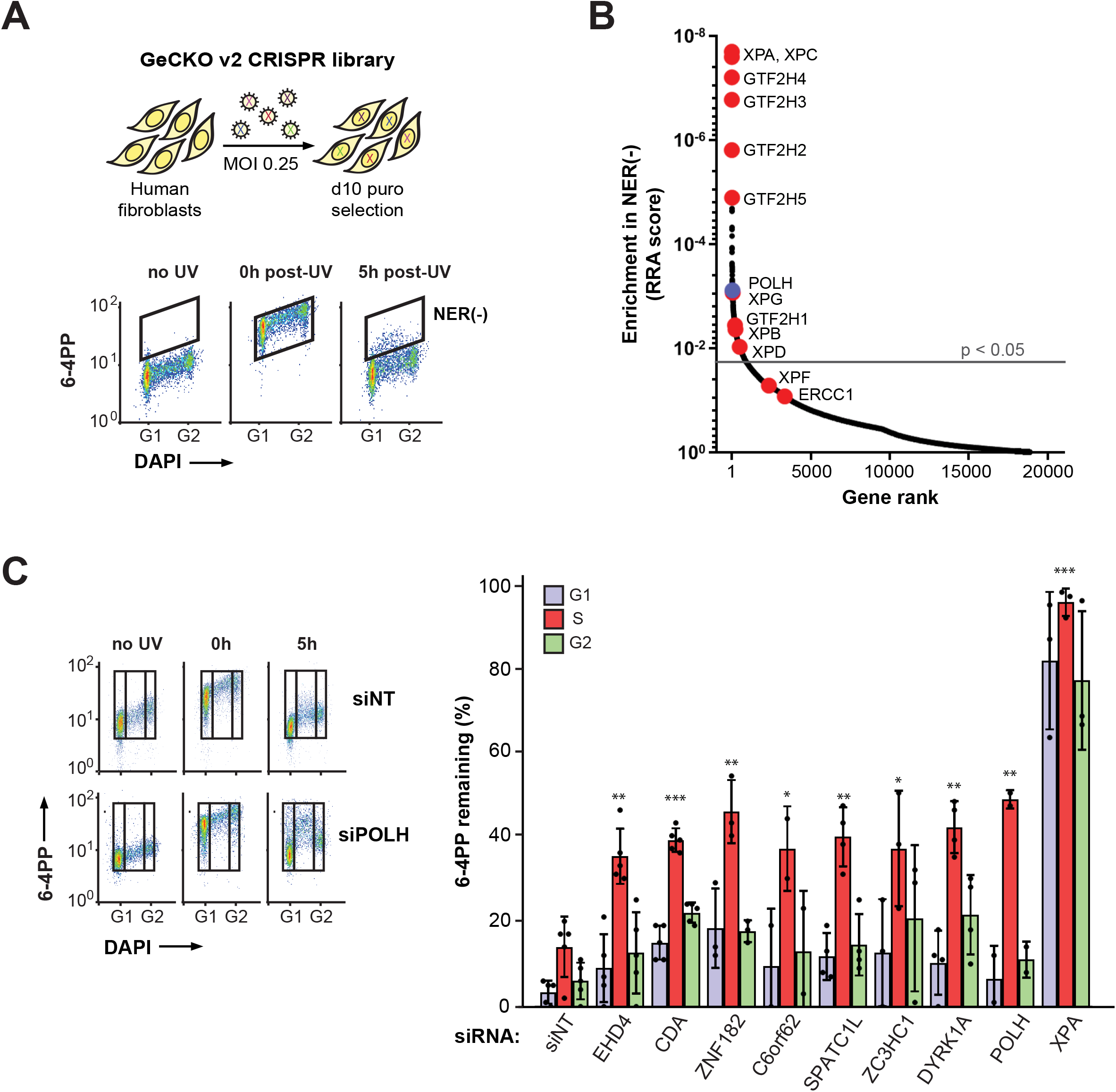
A CRISPR/Cas9 loss-of-function screen identifies novel NER-regulating genes. **A)** LF-1 primary fibroblasts were infected with the GeCKO v2 lentiviral library, irradiated with 20 J/m^2^ of UV, and selected for 10 days in puromycin. A FACS-based NER assay was then employed to sort cells characterized by reduced efficiency of 6-4PP removal (corresponding to the boxed population at 5h-post UV). **B)** MAGeCK software was used to assign an RRA score and adjusted p value for each gene (see Material and Methods). Known GG-NER genes are shown in red and *POLH* (encoding DNA pol eta) in blue. The gray line delineates p = 0.05. **C)** Candidate genes were knocked down individually in HeLa cells using siRNA. Left: representative bivariate plots of 6-4PP removal as a function of cell cycle for cells treated with siRNA against *POLH* or non-targeting (siNT) control. Right: 6-4PP remaining 5h post-UV as a function of cell cycle for individual candidate genes, as well as for *XPA* and *POLH* (controls) following siRNA knockdown. P-values compare the percentage of damage remaining in S for each siRNA vs siNT control.

sgRNAs were PCR-amplified from control and NER-deficient populations, and sequenced. The data were analyzed using MAGeCK software(20) to identify sgRNAs significantly enriched among NER-defective cells (Figure 1B and Table S1). The top 6 “hits” were direct participants in the NER pathway, i.e., genes encoding XPA, XPC, and four TFIIH core subunits (GTF2H2-5); moreover, several other NER pathway genes, including *XP-B, -D, -G* and *GTF2H1* exhibited significant p-values (<0.05). Importantly, consistent with our previous work showing that functional pol eta is required for efficient SPR(8), sgRNAs targeting *POLH* (encoding pol eta) were significantly enriched in the NER-defective fraction. Gene Ontology (GO-term) analysis was performed on the top-ranking 150 genes. As expected, terms associated with NER, DNA repair, and the DNA damage response displayed significant enrichment (Supplementary Table S2). Moreover, all of the significant non-NER/non-DNA-repair GO-terms were populated with NER genes, e.g., the GO-term “DNA templated transcription initiation” includes genes encoding TFIIH subunits. Overall, the data show that our screening strategy is competent in identifying *bona fide* NER-modulating genes.

Among the top 50 genes, we randomly selected seven for validation using HeLa cells as model system: (i) *SPATC1L* (Spermatogenesis and Centriole-Associated Protein 1 Like), a germ-cell specific factor implicated in spermiogenesis(21), (ii) *CDA* (cytidine deaminase), involved in the pyrimidine salvage pathway(22), (iii) *DYRK1A* (Dual-specificity tyrosine phosphorylation-regulated kinase 1A)(23), (iv) *ZC3HC1* (a.k.a. NIPA; Nuclear Interaction Partner Of ALK), a component of an SCF-type E3 ubiquitin ligase complex regulating the G2-M transition(24), (iv) *EHD4* (EH Domain Containing 4), involved in endosomal transport(25), and (v) *C6orf62* and *ZNF182*, with no established functions. Unexpectedly, siRNA-mediated depletion of each of the above factors engendered significant defects in 6-4PP removal during S phase but not G1 or G2, unlike the situation for the NER pathway protein XPA where repair was defective throughout the cell cycle (Figure 1C; see Table S3 for knockdown efficiencies). We note that, as assessed by DAPI staining, the DNA content of S phase HeLa cells did not noticeably increase for at least 5h post-UV, i.e., progression through S was delayed as a result of UV irradiation (Figure S1A-D; the nucleoside analog EdU was used to label S phase cells). Moreover, cells in G1 (EdU-negative) did not enter S during the same period. This likely reflects DNA damage-induced inhibition of the G1-S transition and slow progression through S, and confirms that our experimental conditions permit quantification of 6-4PP removal specifically during S phase, i.e., without any contribution from cells irradiated in G1 which might have then progressed to S. Overall, the data indicate that the human genome encodes previously unknown NER regulators that promote UV DNA photoproduct removal in an S phase-specific manner.

### Dyrk1A promotes NER specifically during S phase and cell survival post-UV

We chose to further characterize the SPR defect in cells depleted for Dyrk1A, a dual specificity kinase that autophosphorylates on a tyrosine residue, but whose numerous substrates are modified on serines and/or threonines. Dyrk1A exhibits broad functionality, being involved in multiple processes associated with neuronal development, transcriptional control, and cell proliferation(26). As was the case for siRNA-mediated knockdown of Dyrk1A in HeLa cells, CRISPR-Cas9 knockout of *Dyrk1A* in primary LF-1 lung fibroblasts significantly inhibited 6-4PP removal uniquely during S phase (Figure 2A), demonstrating that this effect is not cell type-specific. Moreover, quantification of unscheduled incorporation of the nucleoside analog EdU post-UV in G1/G2 HeLa cells did not reveal any effect of Dyrk1A knockdown on NER gap-filling outside of S phase, whereas control cells depleted for the essential NER factor XPA manifested a profound defect (Figure 2B). We further found that knockdown of Dyrk1A sensitized HeLa cells to UV but did not exacerbate the UV sensitivity caused by knockdown of XPA (Figure 2C). This indicates that Dyrk1A protects cells against UV-induced cell killing specifically by promoting NER during S phase.

**FIGURE 2:**
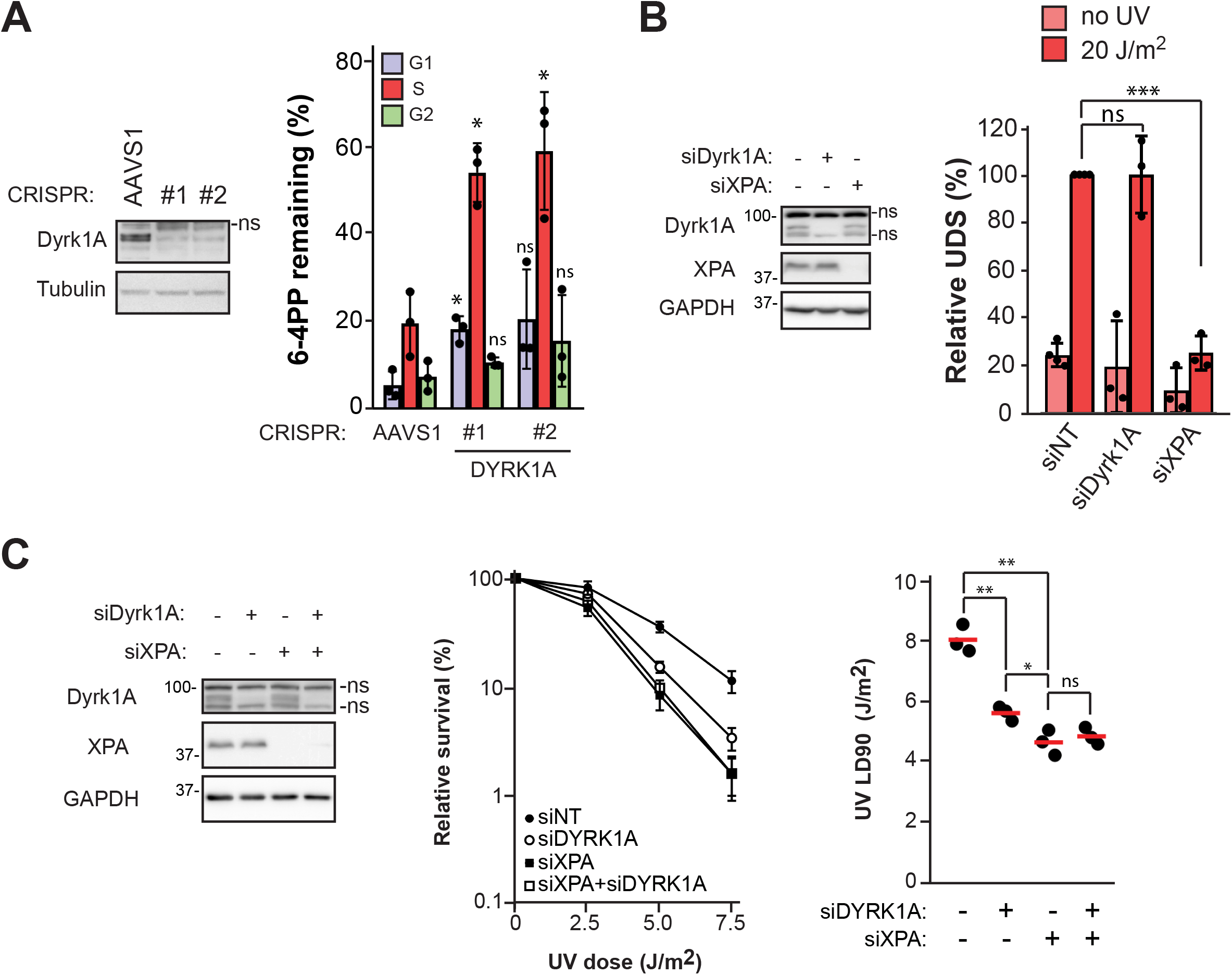
Influence of Dyrk1A on NER and cell survival post-UV. **A)** Left: knockout of *Dyrk1A* by CRISPR-Cas9 in LF-1 primary fibroblasts. sgRNA against the Adeno-Associated Virus Integration Site (AAVS1) was used as negative control. ns on this and all subsequent immunoblots indicates a non-specific band; Right: quantification of 6-4PP removal as in Figure 1C. **B)** Evaluation of unscheduled DNA synthesis (UDS) post-UV in HeLa cells treated with siRNA against Dyrk1A (siDyrk1A) or XPA (siXPA). Left: Immunoblots showing knock-down of Dyrk1A and XPA. Right: Quantification of EdU incorporation in G1/G2 after 20 J/m^2^ UV or mock-treatment. **C)** Clonogenic survival post-UV in HeLa cells treated with siDyrk1A and/or siXPA. Left: immunoblot showing protein knock-down. Middle: clonogenic survival. Right: LD90 values were determined from clonogenic survival curves using GraphPad Prism v8.

### Dyrk1A depletion does not inhibit SPR by causing DNA replication stress

As mentioned earlier, SPR defects can be caused by excessive sequestration of RPA on ssDNA generated during periods of severe replicative stress (10, 11, 14). We therefore sought to examine the impact of Dyrk1A knockdown on the UV-induced replicative stress response. Phosphorylation of histone H2AX(S139), Chk1(S345) and RPA32(S33), well-known markers for activation of replicative stress-induced signaling, were not elevated post-UV in Dyrk1A-depleted vs control HeLa cells (Figure 3A-B). Interestingly, the level of DNA-associated RPA32, which is clearly expected to increase during replicative stress(27), was actually reduced upon Dyrk1A depletion versus non-targeting siRNA controls after treatment with either UV or the replication-blocking drug hydroxyurea (Figure 3C). This was not due to any effect of Dyrk1A on the total amount (Figure 3D) or nuclear localization (Figure 3E) of RPA subunits. Instead, we found that Dyrk1A depletion caused a reduction in total EdU incorporation during S phase (Figure 3F) but did not impact DNA replication fork progression as assessed by DNA fiber assays (Figure 3G). One plausible explanation for these observations is that lack of Dyrk1A might reduce the number of active DNA replication forks/origins. Consistently, the abundance of chromatin-associated MCM7 (a subunit of the MCM replicative helicase complex), and of the replicative processivity factor PCNA, were significantly decreased in S phase cells lacking Dyrk1A (Figure 3H-I). While the precise reason for this remains to be characterized, we highlight previous data indicating that Dyrk1A depletion reduces the duration of G1(16). This, in turn, would be expected to cause a reduction in origin “licensing” thereby diminishing the number of active replication forks(28). We conclude that defective SPR caused by lack of Dyrk1A is not a consequence of replicative stress-induced reduction in RPA availability.

**FIGURE 3:**
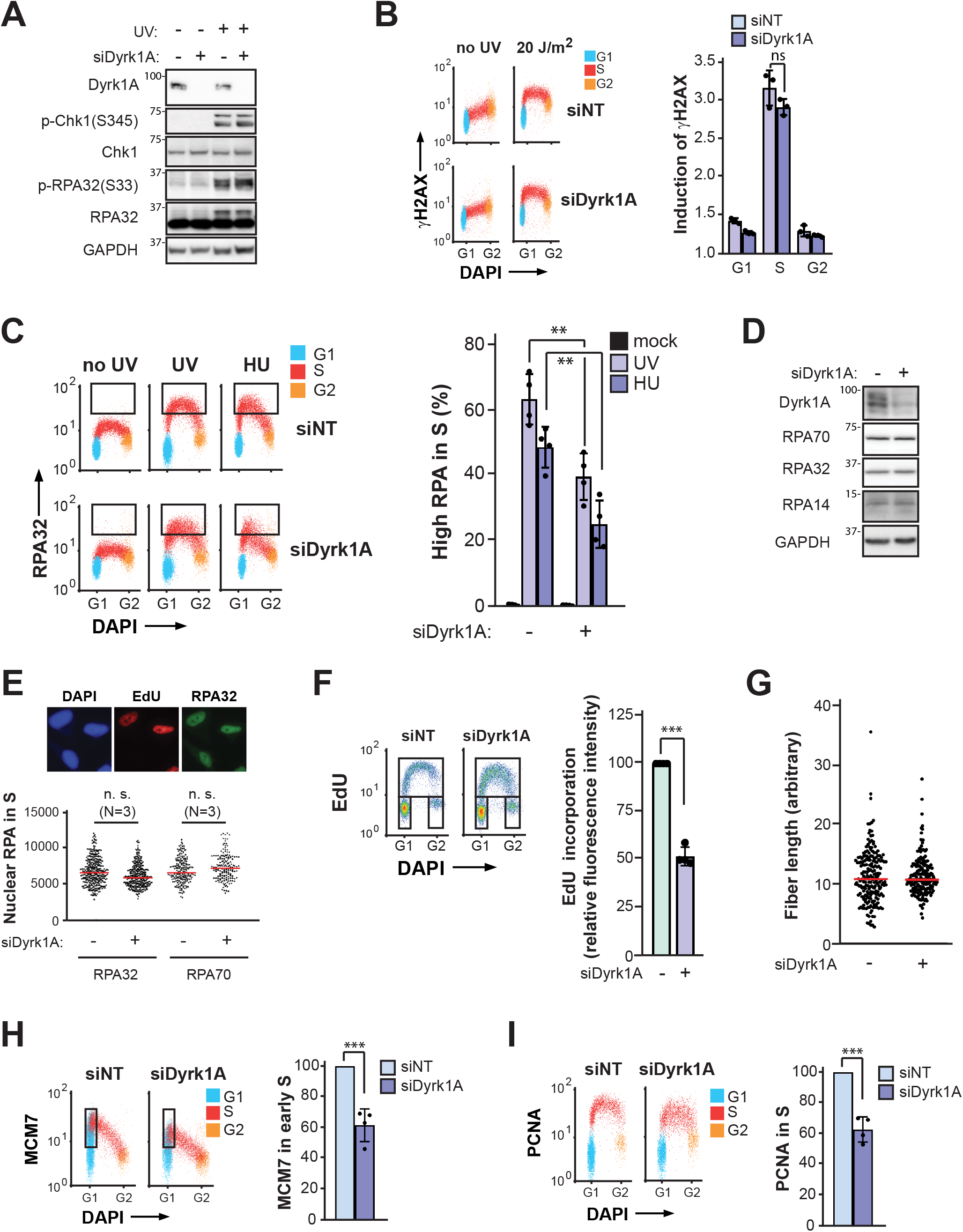
Dyrk1A depletion does not cause replicative stress or reduced RPA availability post-UV. **A)** Detection of phosphorylated Chk1 and RPA32 by immunoblotting 3h after 20 J/m^2^ UV +/- siRNA against Dyrk1A (siDyrk1A) or control (siNT). **B)** Levels of γ-H2AX +/- siDyrk1A were measured as a function of cell cycle 1h after 20 J/m^2^ UV by immunofluorescence flow cytometry. **C)** Left: RPA32-bound DNA was measured +/- siDyrk1A by immunofluorescence flow cytometry 1h after treatment with 20 J/m^2^ UV or 10 mM HU. Cells in various phases of the cell cycle were gated using EdU incorporation as an S phase marker; each phase is represented by a different color. The boxes delineate cells with elevated RPA-bound DNA. Right: quantification of S phase cells with elevated RPA-bound DNA from the left panel. **D)** Immunoblot analysis of RPA subunits from total cellular extracts treated with siDyrk1A or siNT. **E)** Total nuclear levels of RPA32 and RPA70 measured by immunofluorescence. Nuclear intensities for each subunit were quantified from microscopy images. Average intensities of RPA32 or RPA70 in EdU+ cells were determined for siDyrk1A or siNT controls. **F)** Left: Cell cycle distribution assessed by DNA content analysis (DAPI) and EdU incorporation. Right: Fluorescence intensity of EdU incorporation in S phase +/- siDyrk1A. **G)** DNA fiber assay measuring fork progression in HeLa cells +/- siDyrk1A. **H)** Left: MCM7-bound chromatin detected by flow cytometry. Cells in various phases of the cell cycle were gated using EdU incorporation as an S phase marker; each phase is represented by a different color. Right: fluorescence intensity of MCM7-bound chromatin in early S phase (boxed in left panel) for siDyrk1A relative to siNT controls. **I)** Same as H but for PCNA-bound chromatin.

### Dyrk1A depletion causes SPR defects through overexpression of cyclin D1

Among the known substrates of Dyrk1A is cyclin D1, which forms a complex with CDK4/6 to control the G1-S transition(29). Dyrk1A has been shown to phosphorylate cyclin D1 on T286, which targets the latter for timely nuclear export and proteolytic degradation, thus preventing premature entry into S phase(16). Consistently, RNAi knockdown of Dyrk1A in HeLa cells increased cyclin D1 abundance (Figure 4A, left panel), and caused a reduction in the proportion of cells in G1 in a cyclin D1-dependent manner (Figure 4A, right panel). Strikingly, the SPR defect and reduced viability post-UV in cells lacking Dyrk1A were both completely rescued upon co-depletion of cyclin D1 (Figure 4B-C). In contrast, depletion of cyclin D1 did not non-specifically rescue defective SPR caused by ATR inhibition(9) (Figure 4B).

**FIGURE 4:**
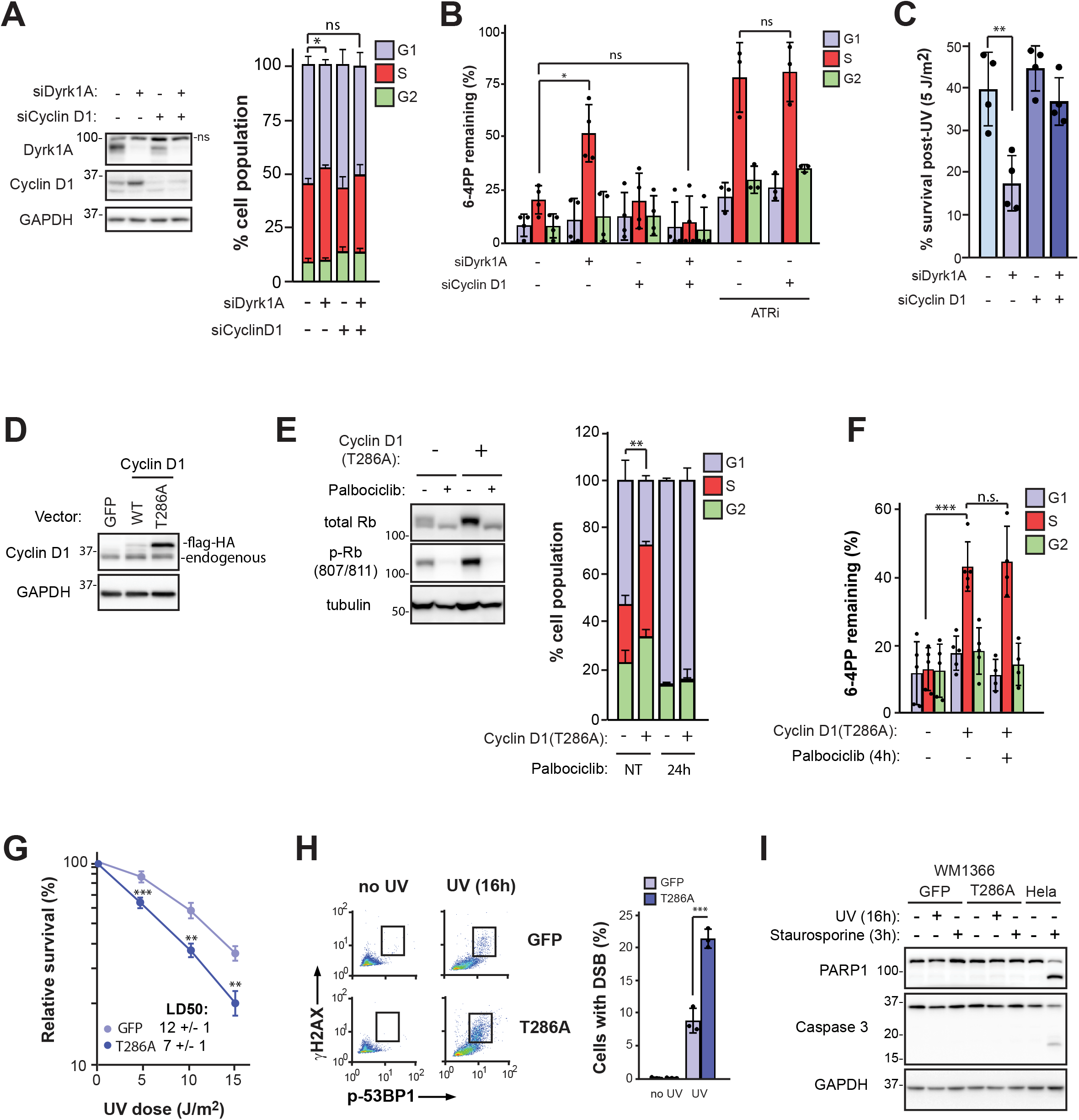
Dyrk1A promotes NER by regulating cyclin D1 stability. **A)** Left: Immunoblot from total HeLa cell extracts following treatment with siRNA against Dyrk1A (siDyrk1A) and/or cyclin D1 (siCyclin D1). Right: cell cycle distribution assessed by DNA content analysis (DAPI) and EdU as in Figure 3F. **B)** Quantification of 6-4PP removal in HeLa cells +/- siDyrk1A and/or siCyclin D1. VE-821 (10 μM) was employed as ATR inhibitor (ATRi). **C)** Clonogenic survival post UV (5 J/m^2^) in HeLa cells +/- siDyrk1A and/or siCyclin D1. **D)** Expression of Flag-HA-cyclin D1 (WT or T286A) in WM1366 using a retroviral construct. Flag-HA-GFP is used as control. **E)** Left: Immunoblot of Rb and phospho-Rb in WM1366 +/- cyclin D1(T286A), pretreated or not for 4 h with 10 μM palbociclib. Right: Overexpression of cyclin D1 (T286A) in WM1366 decreases the % of cells in G1 in a CDK-dependent manner. Cell cycle was assessed by flow cytometry of cells labeled with EdU and DAPI, with or without pre-treatment with the CDK4/6 inhibitor palbociclib for 24h. **F)** Effect of cyclin D1(T286A) overexpression on 6-4PP removal. Cells were pretreated (or not) with palbociclib for 4 h, and the drug was maintained in the medium during post-UV incubation. **G)** UV sensitivity measured by clonogenic survival. **H)** Induction of DSB in WM1366 detected by phospho-53BP1(S1778) and γ-H2AX co-labeling 16h after 20 J/m^2^ UV. **I)** Apoptosis assessed by cleavage of caspase 3 and of PARP1 16h post-UV (1 μM staurosporine for 3 h as positive control).

To further characterize the influence of nuclear cyclin D1 overexpression on NER, we expressed either wild-type (WT) cyclin D1 or the nonphosphorylatable variant cyclin D1(T286A), both epitope-tagged with Flag-HA, in the human melanoma cell line WM1366 which we previously determined is SPR-proficient(30). As expected (16), levels of cyclin D1(T286A)-Flag-HA were much higher compared to the WT counterpart (Figure 4D). The overexpressed T286A variant was functional since it elevated Rb phosphorylation and the fraction of cells in S while concomitantly reducing the proportion of cells in G1; moreover, these phenotypes were reversed by pharmacological inhibition of CDK4/6 using palbociclib (Figure 4E). Importantly, consistent with our result in Dyrk1A-depleted HeLa cells, overexpression of cyclin D1 (T286A) generated an SPR defect in WM1366 (Figure 4F) as well as in LF-1 primary lung fibroblasts (Figure S2A-C). Pre-treatment with palbociclib did not rescue the SPR defect caused by cyclin D1(T286A) expression in WM1366 (Figure 4F), indicating that cyclin D1 regulates SPR in a CDK-independent manner.

Consistent with its negative impact on SPR, cyclin D1(T286A) expression increased UV-induced cytotoxicity (Figure 4G). This was accompanied by elevated DNA double-strand break (DSB) formation at 16 h post-UV, as assessed by induction of histone H2AX(S139) and 53BP1(S1778) phosphorylation (Figure 4H). It appears unlikely that such DSB are due to apoptosis-associated nucleolytic DNA fragmentation(31, 32), since (i) well established markers of apoptosis, i.e., caspase 3- and PARP1-cleavage, were not detected at 16 h after UV irradiation in WM1366 (Figure 4I) and (ii) this cell line appears relatively resistant to apoptosis, as staurosporine treatment did not cause either PARP1- or Caspase 3-cleavage, in contrast to the situation for HeLa cells (Figure 4I). These data indicate that abnormal regulation of cyclin D1, leading to inhibition of NER specifically during S phase, eventually generates lethal DSB and cell death in melanoma cells post-UV.

### Co-stabilization of cyclin D1 and p21 underlies defective SPR in cyclin D1 (T286A)-expressing melanoma cells

Dyrk1A depletion was previously shown to generate enhanced levels of p21 in a cyclin D1-dependent manner(16). Consistently, we observed an increase in p21 abundance upon overexpression of cyclin D1(T286A) in WM1366 cells (Figure 5A); moreover, elevated nuclear expression of both cyclin D1 (T286A) and p21 was observed in S phase as well as G1/G2 cells (Figure 5B-C). Remarkably, siRNA-mediated depletion of p21 resulted in complete rescue of the SPR defect caused by cyclin D1(T286A) expression (Figure 5D). On the other hand, using an inducible Tet-ON system, we found that upregulation of p21 alone in WM1366 cells expressing only endogenous WT cyclin D1 did not result in an SPR defect (Figure 5E-F). We conclude that co-stabilization of p21 is required to inhibit NER during S phase in cyclin D1(T286A)-expressing melanoma cells.

**FIGURE 5:**
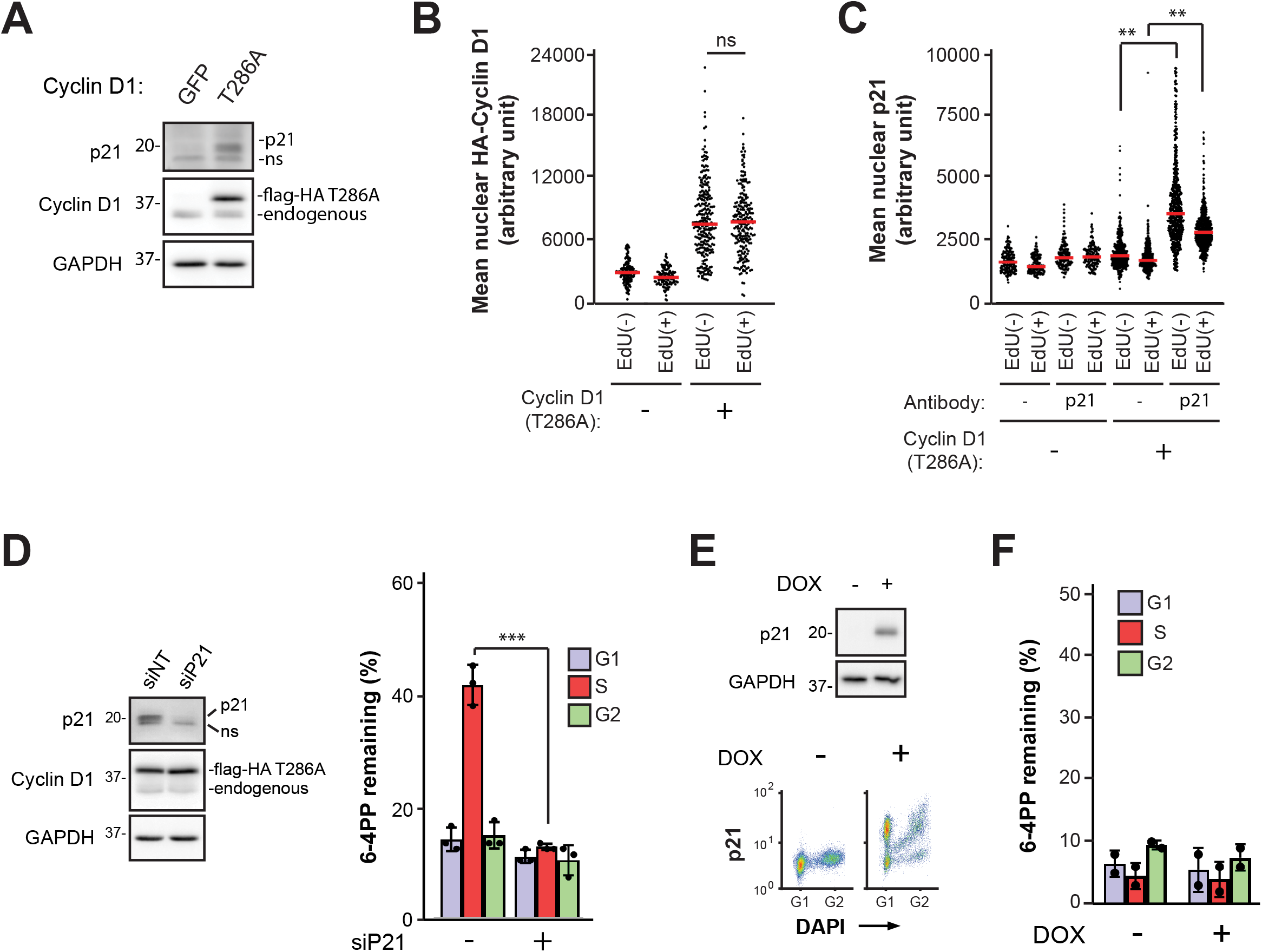
Co-stabilization of p21 is required to sustain the SPR defect caused by cyclin D1 overexpression in melanoma cells. **A)** Immunoblot of p21 and cyclin D1 from total cellular extracts. **B)** Nuclear levels of HA-tagged cyclin D1 in EdU(-) and EdU(+) cells measured by immunofluorescence. **C)** Nuclear levels of p21 in EdU(-) and EdU(+) cells measured by immunofluorescence. **D)** Removal of 6-4PP +/- siRNA against p21 (siP21) in WM1366 overexpressing cyclin D1 (T286A). **E)** Upper: Immunoblot of p21 from WM1366 tet-ON inducible cell line (expressing only endogenous WT cyclin D1) at 5h after induction with doxycycline (DOX). Lower: Cell cycle-specific analysis of p21 protein levels at 5h after induction with DOX. **F)** Removal of 6-4PP at 5h post-UV +/- DOX-induced p21 expression (5h).

## DISCUSSION

We conducted a FACS-based CRISPR/Cas9 screen to identify novel genes implicated in NER. To our knowledge, this represents the first genome-wide functional screen where the efficiency of UV damage repair is the endpoint. While genes encoding NER pathway proteins were well-represented among the top hits, we also recovered *POLH*, encoding pol eta which is known to mediate replicative bypass of UV-induced cyclobutane pyrimidine dimers (33); moreover we previously demonstrated that lack of this translesion DNA polymerase results in abrogation of NER uniquely during S phase(8). The above considerations provide confidence in our screening strategy and show that it can identify genes which influence DNA photoproduct removal in a cell cycle-specific manner.

Remarkably, depletion of all seven genes selected for validation in the present study, which represent cellular pathways with no previously known involvement in NER, generated significant defects in 6-4PP removal strictly in S phase. As discussed earlier, we and others previously demonstrated that such defects can be caused by abnormal replicative stress responses (e.g. in ATR- and pol eta-deficient cells), leading to excessive sequestration of heterotrimeric RPA *in trans* at persistently-stalled replication forks or aberrantly activated origins of replication, which in turn prevents the essential action of this complex in NER(10, 11, 14). Our previously published results demonstrating that SPR defects are (i) recovered in a majority of MM cell lines(30) and (ii) strongly associated with chemosensitivity in ovarian cancer cells(11), highlight the potential biological and clinical relevance of this cell-cycle-specific DNA repair phenotype. Taken together, our findings suggest that a substantial proportion of NER-modulating genes within the human genome, when mutated, cause significant defects in UV damage repair only during S phase. Further investigations are warranted to evaluate whether replicative stress-associated RPA sequestration, and/or other mechanisms, underlie such defects.

We focused our efforts on characterization of the SPR defect caused by lack of Dyrk1A, a kinase which regulates multiple cellular pathways implicated in neurogenesis, transcription, cell cycle control, and cancer development(34). While we initially suspected that Dyrk1A knockdown might reduce SPR efficiency via deregulation of the replicative stress response leading to reduction of RPA availability, our data demonstrate that this is not the case. Rather we found that Dyrk1A regulates SPR through phosphorylation of cyclin D1 on T286(16), a modification that promotes cyclin D1 nuclear export and proteolytic degradation(35, 36). Cyclin D1 forms a complex with CDK4/6 to catalyze phosphorylation of retinoblastoma protein, leading to E2F transcription factor activation and consequent expression of genes required for the G1-S transition(29). As such Dyrk1A-dependent phosphorylation of cyclin D1 contributes to the prevention of premature entry into S phase, and of concomitant genomic instability and enhanced cellular proliferation(16). It should be mentioned that in addition to Dyrk1A, other cellular kinases including GSK3β, ERK1/2, and p38 have been shown to phosphorylate cyclin D1 on T286, which can also promote its cytoplasmic relocalization/proteasomal degradation(37, 38). Nonetheless, in our hands, loss of Dyrk1A alone was sufficient to stabilize and significantly upregulate cyclin D1.

Cyclin D1 is a major driver of multiple cancers(39) including MM(40), through overexpression in either the cytoplasm or nucleus where it promotes tumour invasion/metastasis or deregulation of cell cycle control/enhanced proliferation, respectively. To exert an oncogenic effect in the nucleus, cyclin D1 must be aberrantly stabilized in conjunction with inhibition of its cytoplasmic relocalization(41), as exemplified by the situation for the nonphosphorylatable cyclin D1(T286A) variant used here(35). In addition to its well established role in cell cycle regulation, various non-canonical CDK-dependent and/or -independent roles for nuclear cyclin D1 have been reported, e.g., in transcription and DNA double-strand break repair(42, 43). Moreover, cyclin D1 is known to associate with the essential DNA replication factor PCNA(44–46), thereby forestalling interaction of the latter with DNA polymerases to inhibit DNA synthesis(44). One early investigation indicated that, in UV-exposed human fibroblasts, cyclin D1 overexpression inhibits PCNA-dependent NER gapfilling during G1(47). Our data are not consistent with this, since (i) NER defects caused by cyclin D1 overexpression were observed only in S phase and, (ii) quantification of UDS in HeLa cells revealed no significant impairment of NER gapfilling in either G1 or G2 post-UV upon Dyrk1A knockdown.

p21 (like cyclin D1) is known to interact with PCNA. Indeed p21 strongly binds this DNA replication factor via a canonical PIP box motif(48). However, while there is general agreement that p21 (like cyclin D1) can interfere with DNA replication via PCNA interaction, any influence of this protein on PCNA-dependent gapfilling during NER, or any other step of this repair pathway, remains controversial. Two studies using human cell-free extracts reported that p21 exerts no effect on PCNA-dependent NER gapfilling(49, 50), although another indicated that it strongly inhibits this process(51). Yet another investigation supported this latter result *in vitro*, as well as *in vivo*(52). While our data indicate that Dyrk1A depletion and consequent cyclin D1/p21 overexpression does not negatively impact NER gapfilling during G1 or G2, the possibility of such an impact specifically in S cannot be ruled out. We emphasize that such a possibility would be challenging to evaluate; indeed, quantification by UDS of NER gapfilling during S phase is technically unfeasible, since comparatively weak EdU incorporation signals emanating from this process are dwarfed by those resulting from chromosomal DNA replication(15). While we observed complete rescue of defective SPR in cyclin D1(T286A)-overexpressing melanoma cells upon p21 depletion (without reducing levels of the former), cells expressing only endogenous WT cyclin D1 did not manifest any NER defect following inducible ectopic overexpression of p21. Our overall results thus indicate that co-overexpression of cyclin D1 and p21 is essential to generate the SPR defect caused by Dyrk1A depletion. Further investigation will be necessary to identify the precise underlying mechanism.

Overwhelming evidence demonstrates that efficient removal of UV-induced DNA photoproducts during sunlight exposure is critical for protection against skin cancer. Nonetheless, to date, few data support the plausible expectation that sporadic MM in the general population is frequently characterized by defective NER. We posit that this apparent paradox may be explained, in part, by the fact that no previous studies to our knowledge (with the sole exception of our own revealing a prevalence of SPR defects among model MM cell lines(30)), have considered the possibility that NER might be frequently inhibited in MM in a cell cycle-specific manner. As such, our data showing that aberrant accumulation of nuclear cyclin D1 generates defects in NER specifically during S may harbour major implications for MM development. Moreover, this may extend to other major cancers; for example, carcinoma of the lung, often characterized by expression of oncogenic cyclin D1, is strongly promoted by exposure to the ubiquitous environmental carcinogen benzo(a)pyrene that generates highly-mutagenic DNA adducts repaired exclusively by NER.

## EXPERIMENTAL PROCEDURES

### Cell culture

LF-1 primary human lung fibroblasts(18), a gift from John Sedivy (Brown University), were grown in Eagle’s MEM containing 15% FBS, essential and nonessential amino acids, vitamins and antibiotics (Life Technologies, Carlsbad, CA, USA). HeLa cells (ATCC; Manassas, VA, USA) were cultured in DMEM + 10% FBS and antibiotics. WM1366 melanoma cells (Coriell Institute; Camden, NJ, USA) were cultured as described(30). All cell lines were authenticated by STR analysis (McGill University Genome Center, Montreal, Canada), and routinely tested for mycoplasma contamination by staining with DAPI.

### Reagents and plasmids

Chemical inhibitors, antibodies, siRNAs, and DNA oligonucleotides used in this study are listed in Supporting Experimental Procedures. Plasmids and siRNAs were transfected using Lipofectamine 2000 and RNAiMax, respectively (Life Technologies). LentiCRISPRv2(19) was a gift from Feng Zhang (Addgene plasmid # 52961). cDNAs encoding cyclin D1 WT and cyclin D1 (T286A) were PCR amplified from pcDNA cyclinD1 HA and pcDNA cyclinD1 HA T286A (Addgene plasmids #11181 and # 11182, respectively; gifts from Bruce Zetter (53)), and cloned into pDONR221 using Gateway BP Clonase (Life Technologies). Entry clones were recombined with MSCV-N-Flag-HA-IRES-PURO(54) (a gift from Wade Harper, Addgene plasmid # 41033) using Gateway LR Clonase II (Life Technologies). The coding sequence of p21 was PCR amplified from flag-p21-WT(55) (a gift from Mien-Chie Hung, Abcam #16240) and cloned into pRetro-X-tight-PUR (Clontech), using NotI and EcoRI sites. To generate WM1366-TetON-p21, WM1366 was transduced with pLenti-CMV-rtTA3-Blast (a gift from Eric Campeau, Addgene #26429) and selected with 20 μg/mL blasticidin (Life Technologies). The cell line was then transduced with pRetro-X-tight-PUR-p21 and selected with 1 μg/mL puromycin (Life Technologies). Induction of p21 was carried out with 5 μg/mL of doxycycline (Bioshop Canada, Burlington, Canada) for indicated times.

### Cell irradiation

Cell monolayers were washed with PBS and covered with a thin layer of PBS, followed by irradiation with monochromatic 254-nm UV (hereafter UV) using a G25T8 germicidal lamp (Philips, Cambridge, MA, USA). The fluence was 0.7 J/m^2^/s, as measured with a Spectroline DRC 100x digital radiometer equipped with a DIX-254 sensor (Spectronics Corporation, Melville, NY, USA).

### Flow cytometry–based NER assay

Repair of 6-4PP was evaluated as a function of cell cycle as described(9). Briefly, replicate exponentially-growing cultures were irradiated with 20 J/m^2^ of UV (or mock-irradiated) and harvested either immediately (0h time point) or following 5h incubation to allow repair. Cells were then fixed, permeabilized, and double-stained with DAPI and Alexa647-conjugated anti-6-4PP antibody. Bivariate flow cytometry analysis was used to quantify 6-4PP removal for populations gated in each phase of the cell cycle. Data were acquired using an LSR II flow cytometer (BD Biosciences, Franklin Lakes, NJ, USA) and analyzed with FlowJo software v10 (Ashland, OR, USA).

### CRISPR screen

The GeCKOv2 human CRISPR knockout pooled library (a gift from Feng Zhang, Addgene #1000000048) was used as described (19) to infect a population of LF-1 primary fibroblasts. Following puromycin selection at day 10, 10^8^ sgRNA-transduced cells were irradiated with 20 J/m^2^ UV, labeled with DAPI and anti-6-4PP antibody at 5h post-irradiation, and NER-deficient cells sorted using flow cytometry. Genomic DNA was isolated, and a sample was stained with PicoGreen (Life Technologies) followed by quantification using a TBS-380 fluorometer (Turner Biosystems, Sunnyvale, CA, USA). PCR of genomic DNA and next generation sequencing were carried out as described(19) to identify sgRNAs from control vs NER-deficient populations (see Supporting Experimental Procedures for details). Sequencing adaptors were removed using cutadapt(56). MAGeCK software version 0.5.6(20) was used to generate sgRNA read count data, enriched guides, and gene-level rankings. GO term enrichment was determined with the preranked tool from the GSEA desktop app version 4.1.0 and the Molecular Signatures Database version 7.2(57, 58), using the -log10(P-value) of the top 150 genes obtained by MAGeCK analysis.

### Clonogenic survival

Clonogenic survival post-UV was evaluated as described(30). LD90 values for individual survival curves were determined using GraphPad Prism v8.

### Immunoblotting

Immunoblotting was performed on whole cell extracts using standard protocols. Antibodies are listed in Supporting Experimental Procedures. Imaging was performed using ECL prime reagent (GE healthcare, Chicago, IL, USA) with an Azure c600 instrument (Azure Biosystems, Dublin, CA, USA).

### Cell cycle analysis

Cells were labeled with EdU and DAPI, and analyzed by flow cytometry as described(59). Data were acquired using an LSR II flow cytometer (BD Biosciences) and analyzed with FlowJo software v10.

### Detection of proteins by flow cytometry

Chromatin-bound RPA subunits were detected by flow cytometry as described(59). In the case of PCNA and MCM7, the same protocol was used with the following modifications: After fixation, cells were resuspended in 0.5 mL PBSB (PBS + 0.1% BSA), mixed with 3 mL of −20 ºC methanol and incubated at −20 ºC for 15 min. Cells were pelleted at 1500 rpm for 2 min at 4 °C, washed with 2 mL of PBSB, and washed again with 0.5 mL of 1X BD Perm/Wash buffer (BD Biosciences) before antibody labeling. Detection of p21 was carried out as above except that cells were fixed with 4% formaldehyde in PBS and then permeabilized in cold PBS containing 0.1% triton X100 and 0.3M sucrose on ice for 10 min. In all cases, data were acquired using an LSR II flow cytometer (BD Biosciences) and analyzed using FlowJo software v10.

### Immunofluorescence microscopy

For detection of total RPA32, RPA70, and HA-cyclin D1, cells were pulsed with 10 μM EdU for 20 min, washed with PBS, and fixed with 4% formaldehyde in PBS for 30 min at room temperature. Cells were then permeabilized in 0.1% triton X100 in PBS for 10 min, followed by blocking in PBS containing 3% BSA overnight at 4 ºC. Primary antibodies were added for 3h at room temperature. EdU was then conjugated to Alexa 647 using click-iT chemistry as described(59), followed by DAPI staining. Detection of p21 was performed as above except that 0.2% Triton X-100 was used and blocking was at room temperature in PBS + 3% goat serum + 0.05% Tween-20 for 1 h. Primary antibody was added overnight at 4 ºC. Images were acquired with a DeltaVision Elite system (GE Healthcare). Nuclear fluorescence signals were quantified using custom software as described(10).

### DNA fiber assays

DNA fiber assays were performed as described (60). Images were acquired with a DeltaVision Elite system and individual fibers measured using ImageJ software.

### Unscheduled DNA synthesis

NER gap filling was evaluated by unscheduled DNA synthesis assay using fluorescence microscopy as described(61). Briefly, cells were irradiated with 20 J/m^2^ UV (or mock irradiated) and then incubated for 2h in DMEM without serum, containing 10 μM EdU and 1 μM 5-fluoro-2’-deoxyuridine (Sigma-Aldrich). Images were acquired with a DeltaVision Elite system and nuclear fluorescence signals quantified using custom software as described(10). The relative UDS (%) for each sample was calculated as the median intensity of nuclear EdU divided by that of the UV-treated siNT control. Cells in S-phase, i.e., with saturated EdU signal, were excluded from the analysis.

### Statistical analysis

All experiments were performed independently at least three times (biological replicates), unless stated otherwise. Data are reported as the mean ± SD, except for quantification of microscopy data where medians are shown. Significance was determined with two-tailed unpaired student t-test and adjusted for multiple tests (Holm-Sidak method) where applicable. Statistical analyses were performed using GraphPad Prism v8; p-values are as follows: (*) p-value < 0.05, (**) p-value < 0.01, (***) p-value < 0.001, (ns) p-value not significant.

## Data Availability

Data is available from the corresponding author upon request.

## Supporting Information

This article contains supporting information.

## REFERENCES

1. Brash, D. E., Rudolph, J. A., Simon, J. A., Lin, A., McKenna, G. J., Baden, H. P., et al. (1991) A role for sunlight in skin cancer: UV-induced p53 mutations in squamous cell carcinoma. Proc. Natl. Acad. Sci.. 88, 10124–10128

2. Hodis, E., Watson, I. R., Kryukov, G. V., Arold, S. T., Imielinski, M., Theurillat, J.-P., et al. (2012) A Landscape of Driver Mutations in Melanoma. Cell. 150, 251–263

3. Marteijn, J. A., Lans, H., Vermeulen, W., and Hoeijmakers, J. H. J. (2014) Understanding nucleotide excision repair and its roles in cancer and ageing. Nat. Rev. Mol. Cell Biol. 15, 465–481

4. Lans, H., Hoeijmakers, J. H. J., Vermeulen, W., and Marteijn, J. A. (2019) The DNA damage response to transcription stress. Nature Reviews Molecular Cell Biology. 20, 766–784

5. DiGiovanna, J. J., and Kraemer, K. H. (2012) Shining a light on xeroderma pigmentosum. J. Invest. Dermatol. 132, 785–796

6. Akbani, R., Akdemir, K. C., Aksoy, B. A., Albert, M., Ally, A., Amin, S. B., et al. (2015) Genomic Classification of Cutaneous Melanoma. Cell. 161, 1681–1696

7. Berger, M. F., Hodis, E., Heffernan, T. P., Deribe, Y. L., Lawrence, M. S., Protopopov, A., et al. (2012) Melanoma genome sequencing reveals frequent PREX2 mutations. Nature. 485, 502–506

8. Auclair, Y., Rouget, R., Belisle, J. M., Costantino, S., and Drobetsky, E. A. (2010) Requirement for functional DNA polymerase eta in genome-wide repair of UV-induced DNA damage during S phase. DNA Repair. 9, 754–764

9. Auclair, Y., Rouget, R., Affar, E. B., and Drobetsky, E. A. (2008) ATR kinase is required for global genomic nucleotide excision repair exclusively during S phase in human cells. Proc. Natl. Acad. Sci. 105, 17896–17901

10. Bélanger, F., Angers, J.-P., Fortier, E., Hammond-Martel, I., Costantino, S., Drobetsky, E., et al. (2016) Mutations in Replicative Stress Response Pathways Are Associated with S Phase-Specific Defects in Nucleotide Excision Repair. J. Biol. Chem. 291, 522–37

11. Bélanger, F., Fortier, E., Dubé, M., Lemay, J.-F., Buisson, R., Masson, J.-Y. et al., (2018) Replication Protein A Availability during DNA Replication Stress Is a Major Determinant of Cisplatin Resistance in Ovarian Cancer Cells. Cancer Res. 78, 5561–5573

12. Maréchal, A., and Zou, L. (2015) RPA-coated single-stranded DNA as a platform for post-translational modifications in the DNA damage response. Cell Res. 25, 9–23

13. He, Z., Henricksen, L. A., Wold, M. S., and Ingles, C. J. (1995) RPA involvement in the damage-recognition and incision steps of nucleotide excision repair. Nature. 374, 566–569

14. Tsaalbi-Shtylik, A., Moser, J., Mullenders, L. H. F., Jansen, J. G., and de Wind, N. (2014) Persistently stalled replication forks inhibit nucleotide excision repair in trans by sequestering Replication protein A. Nucleic Acids Res. 42, 4406–4413

15. Kelly, C. M., and Latimer, J. J. (2005) Unscheduled DNA Synthesis. Methods Mol Biol. 291, 303–320

16. Chen, J.-Y., Lin, J.-R., Tsai, F.-C., and Meyer, T. (2013) Dosage of Dyrk1a shifts cells within a p21-cyclin D1 signaling map to control the decision to enter the cell cycle. Mol. Cell. 52, 87–100

17. Musgrove, E. A., Caldon, C. E., Barraclough, J., Stone, A., and Sutherland, R. L. (2011) Cyclin D as a therapeutic target in cancer. Nat. Rev. Cancer. 11, 558–572

18. Brown, J. P., Wei, W., and Sedivy, J. M. (1997) Bypass of senescence after disruption of p21CIP1/WAF1 gene in normal diploid human fibroblasts. Science. 277, 831–834

19. Sanjana, N. E., Shalem, O., and Zhang, F. (2014) Improved vectors and genome-wide libraries for CRISPR screening. Nat. Methods. 11, 783–784

20. Li, W., Xu, H., Xiao, T., Cong, L., Love, M. I., Zhang, F., et al. (2014) MAGeCK enables robust identification of essential genes from genome-scale CRISPR/Cas9 knockout screens. Genome Biol. 15, 554

21. Kim, J., Kwon, J. T., Jeong, J., Kim, J., Hong, S. H., Kim, J., et al. (2018) SPATC1L maintains the integrity of the sperm head-tail junction. EMBO Rep. 19, e45991

22. Frances, A., and Cordelier, P. (2020) The Emerging Role of Cytidine Deaminase in Human Diseases: A New Opportunity for Therapy? Mol Ther. 28, 357–366

23. Laham, A. J., Saber-Ayad, M., and El-Awady, R. (2021) DYRK1A: a down syndrome-related dual protein kinase with a versatile role in tumorigenesis. Cell Mol Life Sci. 78, 603–619

24. Bassermann, F., von Klitzing, C., Münch, S., Bai, R.-Y., Kawaguchi, H., Morris, S. W., et al. (2005) NIPA defines an SCF-type mammalian E3 ligase that regulates mitotic entry. Cell. 122, 45–57

25. Sharma, M., Naslavsky, N., and Caplan, S. (2008) A role for EHD4 in the regulation of early endosomal transport. Traffic. 9, 995–1018

26. Fernández-Martínez, P., Zahonero, C., and Sánchez-Gómez, P. (2015) DYRK1A: the double-edged kinase as a protagonist in cell growth and tumorigenesis. Mol Cell Oncol. 2, e970048

27. Toledo, L. I., Altmeyer, M., Rask, M.-B., Lukas, C., Larsen, D. H., Povlsen, L. K., et al. (2013) ATR prohibits replication catastrophe by preventing global exhaustion of RPA. Cell. 155, 1088–1103

28. Matson, J. P., Dumitru, R., Coryell, P., Baxley, R. M., Chen, W., Twaroski, K., et al. (2017) Rapid DNA replication origin licensing protects stem cell pluripotency. Elife. 6, e30473

29. Sherr, C. J. (1995) D-type cyclins. Trends Biochem Sci. 20, 187–190

30. Bélanger, F., Rajotte, V., and Drobetsky, E. A. (2014) A majority of human melanoma cell lines exhibits an S phase-specific defect in excision of UV-induced DNA photoproducts. PLoS ONE. 9, e85294

31. Panzarino, N. J., Krais, J. J., Cong, K., Peng, M., Mosqueda, M., Nayak, S. U., et al. (2021) Replication Gaps Underlie BRCA Deficiency and Therapy Response. Cancer Res. 81, 1388–1397

32. Rogakou, E. P., Nieves-Neira, W., Boon, C., Pommier, Y., and Bonner, W. M. (2000) Initiation of DNA fragmentation during apoptosis induces phosphorylation of H2AX histone at serine 139. J Biol Chem. 275, 9390–9395

33. Sale, J. E., Lehmann, A. R., and Woodgate, R. (2012) Y-family DNA polymerases and their role in tolerance of cellular DNA damage. Nat. Rev. Mol. Cell Biol. 13, 141–152

34. Rammohan, M., Harris, E., Bhansali, R. S., Zhao, E., Li, L. S., and Crispino, J. D. (2022) The chromosome 21 kinase DYRK1A: emerging roles in cancer biology and potential as a therapeutic target. Oncogene. 10.1038/s41388-022-02245-6

35. Alt, J. R., Cleveland, J. L., Hannink, M., and Diehl, J. A. (2000) Phosphorylation-dependent regulation of cyclin D1 nuclear export and cyclin D1-dependent cellular transformation. Genes Dev. 14, 3102–3114

36. Lin, D. I., Barbash, O., Kumar, K. G. S., Weber, J. D., Harper, J. W. Klein-Szanto, et al. (2006) Phosphorylation-dependent ubiquitination of cyclin D1 by the SCF(FBX4-alphaB crystallin) complex. Mol Cell. 24, 355–366

37. Diehl, J. A., Cheng, M., Roussel, M. F., and Sherr, C. J. (1998) Glycogen synthase kinase-3beta regulates cyclin D1 proteolysis and subcellular localization. Genes Dev. 12, 3499–3511

38. Densham, R. M., Todd, D. E., Balmanno, K., and Cook, S. J. (2008) ERK1/2 and p38 cooperate to delay progression through G1 by promoting cyclin D1 protein turnover. Cell Signal. 20, 1986–1994

39. Tchakarska, G., and Sola, B. (2020) The double dealing of cyclin D1. Cell Cycle. 19, 163–178

40. González-Ruiz, L., González-Moles, M. Á., González-Ruiz, I., Ruiz-Ávila, I., Ayén, Á., and Ramos-García, P. (2020) An update on the implications of cyclin D1 in melanomas. Pigment Cell Melanoma Res. 33, 788–805

41. Kim, J. K., and Diehl, J. A. (2009) Nuclear cyclin D1: an oncogenic driver in human cancer. J Cell Physiol. 220, 292–296

42. Bienvenu, F., Jirawatnotai, S., Elias, J. E., Meyer, C. A., Mizeracka, K., Marson, A., et al. (2010) Transcriptional role of cyclin D1 in development revealed by a genetic-proteomic screen. Nature. 463, 374–378

43. Jirawatnotai, S., Hu, Y., Michowski, W., Elias, J. E., Becks, L., Bienvenu, F., et al. (2011) A function for cyclin D1 in DNA repair uncovered by protein interactome analyses in human cancers. Nature. 474, 230–234

44. Fukami-Kobayashi, J., and Mitsui, Y. (1999) Cyclin D1 inhibits cell proliferation through binding to PCNA and cdk2. Exp Cell Res. 246, 338–347

45. Matsuoka, S., Yamaguchi, M., and Matsukage, A. (1994) D-type cyclin-binding regions of proliferating cell nuclear antigen. J Biol Chem. 269, 11030–11036

46. Xiong, Y., Zhang, H., and Beach, D. (1992) D type cyclins associate with multiple protein kinases and the DNA replication and repair factor PCNA. Cell. 71, 505–514

47. Pagano, M., Theodoras, A. M., Tam, S. W., and Draetta, G. F. (1994) Cyclin D1-mediated inhibition of repair and replicative DNA synthesis in human fibroblasts. Genes Dev. 8, 1627–1639

48. Mansilla, S. F., de la Vega, M. B., Calzetta, N. L., Siri, S. O., and Gottifredi, V. (2020) CDK-Independent and PCNA-Dependent Functions of p21 in DNA Replication. Genes (Basel). 11, E593

49. Shivji, M. K., Grey, S. J., Strausfeld, U. P., Wood, R. D., and Blow, J. J. (1994) Cip1 inhibits DNA replication but not PCNA-dependent nucleotide excision-repair. Curr Biol. 4, 1062–1068

50. Li, R., Waga, S., Hannon, G. J., Beach, D., and Stillman, B. (1994) Differential effects by the p21 CDK inhibitor on PCNA-dependent DNA replication and repair. Nature. 371, 534–537

51. Pan, Z. Q., Reardon, J. T., Li, L., Flores-Rozas, H., Legerski, R., Sancar, A., and Hurwitz, J. (1995) Inhibition of nucleotide excision repair by the cyclin-dependent kinase inhibitor p21. J Biol Chem. 270, 22008–22016

52. Cooper, M. P., Balajee, A. S., and Bohr, V. A. (1999) The C-terminal domain of p21 inhibits nucleotide excision repair In vitro and In vivo. Mol Biol Cell. 10, 2119–2129

53. Newman, R. M., Mobascher, A., Mangold, U., Koike, C., Diah, S., Schmidt, M., et al. (2004) Antizyme targets cyclin D1 for degradation. A novel mechanism for cell growth repression. J Biol Chem. 279, 41504–41511

54. Sowa, M. E., Bennett, E. J., Gygi, S. P., and Harper, J. W. (2009) Defining the human deubiquitinating enzyme interaction landscape. Cell. 138, 389–403

55. Zhou, B. P., Liao, Y., Xia, W., Spohn, B., Lee, M. H., and Hung, M. C. (2001) Cytoplasmic localization of p21Cip1/WAF1 by Akt-induced phosphorylation in HER-2/neu-overexpressing cells. Nat Cell Biol. 3, 245–252

56. Martin, M. (2011) Cutadapt removes adapter sequences from high-throughput sequencing reads. EMBnet.journal. 17, 10–12

57. Subramanian, A., Tamayo, P., Mootha, V. K., Mukherjee, S., Ebert, B. L., Gillette, M. et al. (2005) Gene set enrichment analysis: a knowledge-based approach for interpreting genome-wide expression profiles. Proc. Natl. Acad. Sci. 102, 15545–15550

58. Liberzon, A., Birger, C., Thorvaldsdóttir, H., Ghandi, M., Mesirov, J. P., and Tamayo, P. (2015) The Molecular Signatures Database (MSigDB) hallmark gene set collection. Cell Syst. 1, 417–425

59. Forment, J. V., and Jackson, S. P. (2015) A flow cytometry-based method to simplify the analysis and quantification of protein association to chromatin in mammalian cells. Nat Protoc. 10, 1297–1307

60. Quinet, A., Carvajal-Maldonado, D., Lemacon, D., and Vindigni, A. (2017) DNA Fiber Analysis: Mind the Gap! Meth. Enzymol. 591, 55–82

61. Limsirichaikul, S., Niimi, A., Fawcett, H., Lehmann, A., Yamashita, S., and Ogi, T. (2009) A rapid non-radioactive technique for measurement of repair synthesis in primary human fibroblasts by incorporation of ethynyl deoxyuridine (EdU). Nucleic Acids Res. 37, e31

