## Supplemental Files for "A genome-wide screen reveals that Dyrk1A kinase promotes nucleotide excision repair by preventing aberrant co-stabilization of cyclin D1 and p21"

**Contains :**

- Figure S1

- Figure S2

- Table S1 (separate xlsx file)

- Table S2 (separate xlsx file)

- Table S3

- Supporting experimental procedures

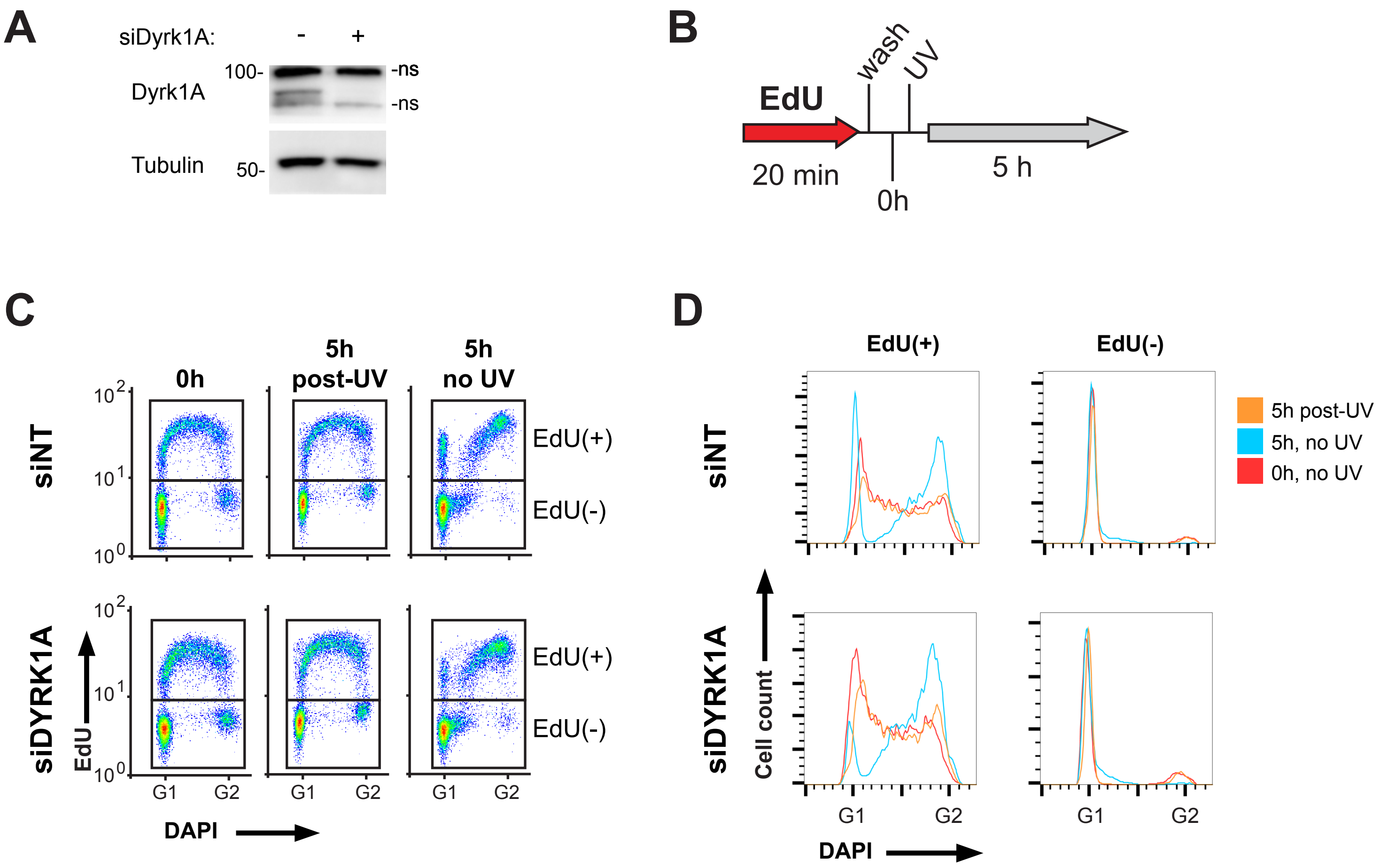


**Figure S1.**

**Progression of HeLa cells through the cell cycle post UV.**

**A)** Western blot showing Dyrk1A levels at 48h post-transfection with siDyrk1A or non-targeting control. **B)** Cells were pulsed with EdU for 20 min and washed with PBS. A sample was collected immediately after the EdU pulse (0h). Samples were irradiated with UV (or mock-irradiated) and harvested 5h later. **C)** Flow cytometry of cells labeled with DAPI and EdU. Boxes show gating of EdU(-) cells, and of EdU(+) cells which were in S at the time of UV irradiation. **D)** Cell cycle distribution of (DAPI-stained) EdU(-) and EdU(+) populations as gated in panel C. In the absence of UV (blue line), EdU(+) cells progressed toward G2/M, with a fraction cycling back into G1. A fraction of EdU(-) cells also entered into S. At 5h post-UV (orange line), there was almost no progression of EdU(+) cells (compare with 0h control; red line). There were also no EdU(-) cells entering into S.

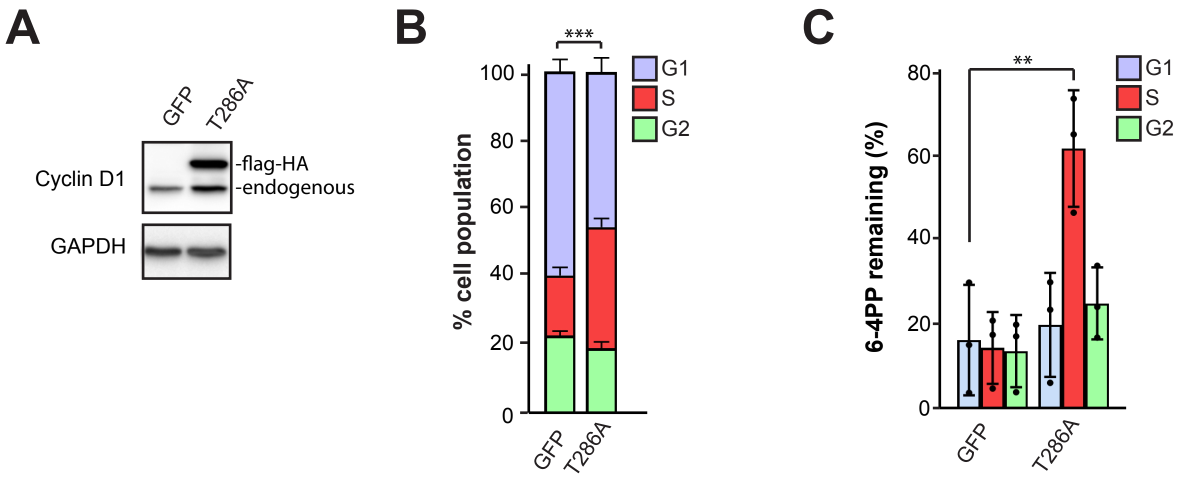


**Figure S2.**
**Stable expression of cyclin D1 (T286A) and negative control (GFP) in LF-1 primary human fibroblasts.**
**A)** Western blot of cyclin D1 using total protein extracts. **B)** Proportion of cells in each phase of the cell cycle. **C)** Removal of 6-4PP at 5h post UV. Values are average +/- SD of 3 independent experiments (unpaired t test: ** $p < 0.01$; ***$p < 0.001$).

**Table S3**
**RT-qPCR quantification of siRNA knock-down efficiencies in Hela cells**

| mRNA | % remaining | Primer pairs used for qPCR | |
|---|---|---|---|
| HPRT | control | FWD | CCCTGGCGTCGTGATTAGTG |
| | | REV | TCGAGCAAGACGTTCAGTCC |
| EHD4 | 10 +/- 2 | FWD | CGCCGTGATGTATGGAGAGA |
| | | REV | GGGAGCTGTGAGCACATGAA |
| SPATC1L | 4 +/- 2 | FWD | CCAGGAGGAGCTACTACCTCAA |
| | | REV | CCACCGTGAAGCCGTAGAG |
| ZNF182 | 23 +/- 1 | FWD | GGCAGCAGGTTACCAAACCA |
| | | REV | GGTCATCCTGATGTTGCCTCT |
| C6ORF62 | 14 +/- 4 | FWD | TCTCTAGCTGACCAGTTTGACT |
| | | REV | TCTCGCACACCTTTCAGGAT |
| DYRK1A | 25 +/- 3 | FWD | TTGTCATGTTACAGAGGCGG |
| | | REV | CCCTGTTGGTGTCTTCGCTT |
| ZC3HC1 | 7 +/- 5 | FWD | ACGCGAAGGACACGTCTG |
| | | REV | AGCTCAAAGGGCTTACCTGC |
| POLH | 40 +/- 5 | FWD | AACCCAGCATTTTCGGCAAC |
| | | REV | TCCATGTCCACGAGAGCAAC |
| CDA | 4 +/- 3 | FWD | CAGTCACTTTCCTGTGGGGG |
| | | REV | GAGACGGCCTTCTGGATAGC |
| XPA | 17 +/- 4 | FWD | GAAGCAAAGGAAGTCCGACAG |
| | | REV | ACACGCTGCTTCTTACTGCT |

RNA was extracted with Trizol reagent (Life Technologies) and cDNA prepared with M-MuLV Reverse Transcriptase (New England Biolabs, Ipswich, MA, USA). Quantitative PCR was performed using Luna Universal qPCR Master Mix (New England Biolabs) on an ABI 7500 instrument (Applied Biosystems; Waltham, Mass, USA). Remaining mRNA (relative to siNT) was calculated by ΔΔCt using HPRT as the housekeeping control. Values are average +/- SD from at least 3 independent experiments

### SUPPORTING EXPERIMENTAL PROCEDURES

**Chemical inhibitors used in this study:**

| Name: | Catalog: | Supplier: |
|---|---|---|
| Palbociclib | PZ0383 | Sigma-Aldrich (Oakville, Canada) |
| PHA-767491 | #18218 | Cayman Chemical Co (Ann Arbor, MI, USA) |
| Staurosporine | # 81590 | Cayman Chemical Co. |
| VE-821 ATR inhibitor | #A2521 | ApexBio (Houston, TX, USA) |
| Hydroxyurea | HYD023 | Bioshop Canada (Burlington, Canada) |

**PCR for Illumina sequencing of Gecko library:**

Genomic DNA was extracted from cell pellets using SNET buffer (20 mM Tris-HCl pH8, 400 mM NaCl, 1% SDS, 5 mM EDTA) with 0.4 mg/ml proteinase K (Life Technologies, Carlsbad, CA, USA) at 55 ºC overnight, followed by extraction with phenol/chloroform (Life Technologies) and treatment with RNase A (BioBasic, Markham, Canada) for 30 minutes at 37 ºC. DNA was quantified with PicoGreen (Life Technologies) using a TBS-380 fluorimeter (Turner Biosystems, Sunnyvale, CA, USA). PCR for Illumina sequencing were prepared using NEBNext High-Fidelity 2X PCR Master Mix, following manufacturer's instructions (New England Biolabs, Ipswich, MA, USA). A first PCR of 20 cycles was prepared in multiple reactions for the total amount of gDNA, at 4 µg DNA per reaction.

PCR 1 Forward: AATGGACTATCATATGCTTACCGTAACTTGAAAGTATTTCG

PCR 1 Reverse: TCTACTATTCTTTCCCCTGCACTGTTGTGGGCGATGTGCGCTCTG

Identical reactions were combined and a second PCR of 24 cycles was performed using 5 µL of PCR1 as template with primers containing Illumina sequences and barcodes (8 identical reactions were done for each sample and combined). PCR2 products were purified on a 1.5% agarose TAE gel, visualized with SYBR-safe (Life Technologies). Sequencing was performed on an Illumina NextSeq system at the Institut de Recherche en Immunologie et Cancérologie (Montréal, Canada).

Primers used for PCR 2:

PCR 2 Forward primers (10 staggered oligos used as a mix):

V2-F0:
AATGATACGGCGACCACCGAGATCTACACTCTTTCCCTACACGACGCTCTTCCGATCTTC
TTGTGGAAAGGACGAAACACCG

V2-F1
AATGATACGGCGACCACCGAGATCTACACTCTTTCCCTACACGACGCTCTTCCGATCTTT
CTTGTGGAAAGGACGAAACACCG

V2-F2
AATGATACGGCGACCACCGAGATCTACACTCTTTCCCTACACGACGCTCTTCCGATCTAT
TCTTGTGGAAAGGACGAAACACCG

V2-F3
AATGATACGGCGACCACCGAGATCTACACTCTTTCCCTACACGACGCTCTTCCGATCTGA
TTCTTGTGGAAAGGACGAAACACCG

V2-F4
AATGATACGGCGACCACCGAGATCTACACTCTTTCCCTACACGACGCTCTTCCGATCTCG
ATTCTTGTGGAAAGGACGAAACACCG

V2-F5
AATGATACGGCGACCACCGAGATCTACACTCTTTCCCTACACGACGCTCTTCCGATCTGC
GATTCTTGTGGAAAGGACGAAACACCG

V2-F6
AATGATACGGCGACCACCGAGATCTACACTCTTTCCCTACACGACGCTCTTCCGATCTAG
CGATTCTTGTGGAAAGGACGAAACACCG

V2-F7
AATGATACGGCGACCACCGAGATCTACACTCTTTCCCTACACGACGCTCTTCCGATCTGAGCGATTCTTGTGGAAAGGACGAAACACCG

V2-F8
AATGATACGGCGACCACCGAGATCTACACTCTTTCCCTACACGACGCTCTTCCGATCTCGAGCGATTCTTGTGGAAAGGACGAAACACCG

V2-F9
AATGATACGGCGACCACCGAGATCTACACTCTTTCCCTACACGACGCTCTTCCGATCTACGAGCGATTCTTGTGGAAAGGACGAAACACCG

PCR2 Reverse primer with Illumina Index #6:

CAAGCAGAAGACGGCATACGAGATGCCAATGTGACTGGAGTTCAGACGTGTGCTCTTCCGATCTTTCTACTATTCTTTCCCTGCACTGT

PCR2 Reverse primer with Illumina Index #12:

CAAGCAGAAGACGGCATACGAGATCTTGTAGTGACTGGAGTTCAGACGTGTGCTCTTCCGATCTTATCTACTATTCTTTCCCCTGCACTGT

**PCR primers for Gateway cloning**

CCND1 Forward:

GGGGACAAGTTTGTACAAAAAAGCAGGCTTCATGGAACACCAGCTCCTGTGCTGC

CCND1 Reverse:

GGGGACCACTTTGTACAAGAAAGCTGGGTCTCAGATGTCCACGTCCCGCACGTC

EGFP Forward:

GGGGACAAGTTTGTACAAAAAAGCAGGCTTCATGGTGAGCAAGGGCGAGGAGCTG

EGFP Reverse:

GGGGACCACTTTGTACAAGAAAGCTGGGTCTTACTTGTACAGCTCGTCCATGCCG

**PCR primers for p21 cloning**

p21 Forward

GGAGCCGCGGCCGCCGCCACCATGTCAGAACCGGCTGGGGATG

p21 Reverse

GAGTCCGAATTCTTAGGGCTTCCTCTTGGAGAAGATC

**DNA cassettes for cloning into pLentiCRISPRv2**

DYRK1A #1 Forward

CACCGTATGACTAGAATCGTCTCCC

DYRK1A #1 Reverse

AAACGGGAGACGATTCTAGTCATAC

DYRK1A #2 Forward

CACCGAAGAAGCGAAGACACCAACA

DYRK1A #2 Reverse

AAACTGTTGGTGTCTTCGCTTCTTC

AAVS1 Forward

CACCGGGGGCCACTAGGGACAGGAT

AAVS1 Reverse

AAACATCCTGTCCCTAGTGGCCCCC

**Antibodies used in this study**

Dilutions of antibodies were 1:1000 for Western blotting (WB), 1:100 for flow cytometry (FC) and 1:300 for immunofluorescence (IF).

| Name: | Catalog: | Supplier: | Applications: |
|---|---|---|---|
| rat anti-Tubulin | ab6161 | Abcam (Cambridge, UK) | WB |
| rabbit anti-RPA1 | ab79398 | Abcam | IF, WB |
| mouse anti-RPA3 | ab167593 | Abcam | WB |
| rabbit anti-Chk1 phospho S345 | #2348 | New England Biolabs | WB |
| rabbit anti-PARP | #9532S | New England Biolabs | WB |
| mouse anti-Rb | #9309 | New England Biolabs | WB |
| rabbit anti-Rb (pS807/811) | #9308 | New England Biolabs | WB |
| mouse anti-Chk1 | sc-8408 | Santa Cruz (Dallas TX, USA) | WB |
| mouse anti-GAPDH | sc-365062 | Santa Cruz | WB |
| mouse anti-caspase 3 | sc-56053 | Santa Cruz | WB |
| mouse anti-PCNA | sc-56 | Santa Cruz | FC |
| rabbit anti-p21 | sc-397 | Santa Cruz | WB |
| mouse anti-p21 | #556430 | BD (Franklin Lakes NJ, USA) | IF, FC |
| rabbit anti-cyclin D1 | GTX108624 | GeneTex (Irvine CA, USA) | WB |
| mouse anti-XPA | GTX72316 | GeneTex | WB |
| rabbit anti-Dyrk1A | #6255 | ProSci (Fort Collins CO, USA) | WB |
| mouse anti-RPA2 | NA18 | Sigma-Aldrich | WB, IF, FC |
| rabbit anti-RPA2 (p-S33) | A300-246A | Bethyl (Montgomery TX, USA) | WB |

| | | | |
|---|---|---|---|
| mouse anti-gH2AX | JBW301 | Sigma-Aldrich | FC |
| rabbit anti-53BP1(p-S1778) | #2675 | Cell Signaling (Danvers MA, USA) | FC |
| mouse anti-MCM7 | sc-9966 | Santa-Cruz | FC |
| mouse anti-HA (12CA5) | ab1424 | Abcam | IF |
| mouse anti-(6-4) photoproducts | MC-082 | Kamiya, Seattle, WA | FC |
| anti-mouse IgG AlexaFluor488 | A11029 | Life Technologies | IF, FC |
| anti-rabbit IgG Alexa Fluor594 | A11012 | Life Technologies | IF, FC |
| anti-rabbit IgG Alexa Fluor488 | A11008 | Life Technologies | IF, FC |
| anti-rabbit IgG-HRP | sc-2357 | Santa Cruz | WB |
| m-IgGk-BP-HRP | sc-516102 | Santa Cruz | WB |
| anti-Rat IgG-HRP | ab97057 | Abcam | WB |

**siRNA oligo duplexes**

Individual duplexes were purchased from Sigma-Aldrich and used as a pooled mix for each gene.

| Gene | Catalog # | Sense sequence: |
|---|---|---|
| DYRK1A | SASI_Hs01_00123259 | GAGCUAUGGACGUUAAUUU[dT][dT] |
| DYRK1A | SASI_Hs01_00123260 | GUGCAAUCAAGAUAGUUGA[dT][dT] |
| DYRK1A | SASI_Hs01_00123261 | GGGUAUUCCACCUGCUCAU[dT][dT] |
| DYRK1A | SASI_Hs01_00123262 | CUCUUUGAACCUAACACGA[dT][dT] |
| CCND1 | SASI_Hs01_00213908 | GCAUGUUCGUGGCCUCUAA[dT][dT] |
| CCND1 | SASI_Hs01_00213909 | CCACAGAUGUGAAGUUCAU[dT][dT] |
| XPA | SASI_Hs02_00302440 | GGAAAGAAUUUAUGGAUUC[dT][dT] |
| XPA | SASI_Hs01_00233870 | CUACUGGAGGCAUGGCUAA[dT][dT] |

| | | |
|---|---|---|
| XPA | SASI_Hs02_00302441 | GGAUUCUUAUCUUAUGAAC[dT][dT] |
| XPA | SASI_Hs01_00233868 | CACUUUGAUUUGCCAACUU[dT][dT] |
| CDA | SASI_Hs01_00229075 | CUGGAGAACUUCAUAAAGA[dT][dT] |
| CDA | SASI_Hs02_00332546 | CCAGUGACAUGCAAGAUGA[dT][dT] |
| CDA | SASI_Hs01_00229077 | GCACCAACUGGCCCGUGUA[dT][dT] |
| CDA | SASI_Hs01_00229078 | CCUACAGGGACUGGGCAAA[dT][dT] |
| EHD4 | SASI_Hs01_00139749 | CAUCUCAGAUGAAUUCUCA[dT][dT] |
| EHD4 | SASI_Hs01_00139750 | CCUUCAUCGCCGUGAUGUA[dT][dT] |
| EHD4 | SASI_Hs01_00139751 | GCUAUAUUUAGAGUGGCAA[dT][dT] |
| EHD4 | SASI_Hs01_00139752 | GGUACUGCGCGUCUACAUU[dT][dT] |
| POLH | SASI_Hs01_00088494 | CAUUGAUGAGGCUUACGUA[dT][dT] |
| POLH | SASI_Hs01_00088495 | CAUAGAGAGGGAGACUGGU[dT][dT] |
| POLH | SASI_Hs01_00088497 | CCAAAUGCCCAUUCGCAAA[dT][dT] |
| POLH | SASI_Hs02_00341708 | GAAUAAACCUUGUGCAGUU[dT][dT] |
| SPATC1L | SASI_Hs01_00105335 | GAGCAGACCUCCACCAAGU[dT][dT] |
| SPATC1L | SASI_Hs01_00105337 | GCGAGUUCCUCAUCAACAC[dT][dT] |
| SPATC1L | SASI_Hs01_00105338 | GGAGCUACUACCUCAAUGA[dT][dT] |
| SPATC1L | SASI_Hs01_00105339 | CUCAUCAACACCUACGGAA[dT][dT] |
| ZC3HC1 | SASI_Hs01_00207894 | CAGAUUGAAUCGUCCAUGA[dT][dT] |
| ZC3HC1 | SASI_Hs01_00207895 | GGAACAACCUUCAUUGGAA[dT][dT] |
| ZC3HC1 | SASI_Hs01_00207896 | CAAGAUCAGUCUUCUCCUA[dT][dT] |
| ZC3HC1 | SASI_Hs01_00207897 | CCUUCAUUGGAAUCUACAA[dT][dT] |
| ZNF182 | SASI_Hs01_00046638 | GCAACAUCAGGAUGACCAA[dT][dT] |

| | | |
|---|---|---|
| ZNF182 | SASI_Hs01_00046639 | GUCAAACCUUGGUGUACAU[dT][dT] |
| ZNF182 | SASI_Hs01_00046640 | CACUGAAAGAAAGUUGUGU[dT][dT] |
| ZNF182 | SASI_Hs01_00046641 | CUAAUCAGCAGUAUCUAUU[dT][dT] |
| C6orf62 | SASI_Hs01_00099792 | GUACUCAGGAGAUGGAUUU[dT][dT] |
| C6orf62 | SASI_Hs01_00099794 | GACAAAUAAUUAUGAAGAA[dT][dT] |
| C6orf62 | SASI_Hs01_00099795 | GUUAUACCAGUCAUGACAA[dT][dT] |
| C6orf62 | SASI_Hs01_00099796 | GAGGUUAUACCAGUCAUGA[dT][dT] |
| P21 | SASI_Hs01_00025255 | CUAAGAGUGCUGGGCAUUU[dT][dT] |
| P21 | SASI_Hs01_00025256 | CUGUCACAGGCGGUUAUGA[dT][dT] |
| P21 | SASI_Hs01_00025257 | CCCUAAUCCGCCCACAGGA[dT][dT] |
